# In silico detection of SARS-CoV-2 specific B-cell epitopes and validation in ELISA for serological diagnosis of COVID-19

**DOI:** 10.1101/2020.05.22.111526

**Authors:** Isabelle Q. Phan, Sandhya Subramanian, David Kim, Lauren Carter, Neil King, Ivan Anishchenko, Lynn K. Barrett, Justin Craig, Logan Tillery, Roger Shek, Whitney E. Harrington, David M. Koelle, Anna Wald, Jim Boonyaratanakornkit, Nina Isoherranen, Alexander L. Greninger, Keith R. Jerome, Helen Chu, Bart Staker, Lance Stewart, Peter J. Myler, Wesley C. Van Voorhis

## Abstract

Rapid generation of diagnostics is paramount to understand epidemiology and to control the spread of emerging infectious diseases such as COVID-19. Computational methods to predict serodiagnostic epitopes that are specific for the pathogen could help accelerate the development of new diagnostics. A systematic survey of 27 SARS-CoV-2 proteins was conducted to assess whether existing B-cell epitope prediction methods, combined with comprehensive mining of sequence databases and structural data, could predict whether a particular protein would be suitable for serodiagnosis. Nine of the predictions were validated with recombinant SARS-CoV-2 proteins in the ELISA format using plasma and sera from patients with SARS-CoV-2 infection, and a further 11 predictions were compared to the recent literature. Results appeared to be in agreement with 12 of the predictions, in disagreement with 3, while a further 5 were deemed inconclusive. We showed that two of our top five candidates, the N-terminal fragment of the nucleoprotein and the receptor-binding domain of the spike protein, have the highest sensitivity and specificity and signal-to-noise ratio for detecting COVID-19 sera/plasma by ELISA. Mixing the two antigens together for coating ELISA plates led to a sensitivity of 94% (N=80 samples from persons with RT-PCR confirmed SARS-CoV2 infection), and a specificity of 97.2% (N=106 control samples).

## Introduction

The COVID-19 pandemic has highlighted the importance of diagnostic tools that are accurate and cost-effective. So far reports based on linear epitope scanning have been limited in scope to a few selected proteins^1^. Recent antibody response results are based on single SARS-CoV-2 proteins^2,3^, or limit testing of the polyprotein to a single domain^4^. We propose a simple method to predict likely candidates for serological diagnostics based on existing prediction tools for B-cell epitopes that leverages available structural and sequencing data.

## Results and Discussion

### Epitope predictions

One of our aims is to propose a visual aid to select suitable targets for the serological diagnosis of a viral pathogen. Results come in the form of a <u>Summary Plot</u> for each viral protein (Suppl **Figure S1**).

First, linear epitopes were predicted on the sequence of each protein with BepiPred2^5^. At the chosen threshold (80% specificity/30% sensitivity)^6^, epitopes were detected for ⅔ of the proteins. Distributions per protein and length show that over half the predicted epitopes were located on only 4 proteins: S, nsp3, N and nsp12 (**Figure 1A**) and that over 80% of the epitopes were below 15 residues long (**Figure 1B**). The raw epitope scores plotted against the sequence illustrates how heavily the threshold affects the prediction (**Figure S1 A**). This is a known limitation of the predictor which achieves high specificity at a high sensitivity cost. Nonetheless, strong signals were observed for the nucleoprotein, consistent with several recent studies^2,7^.

**Figure 1.**
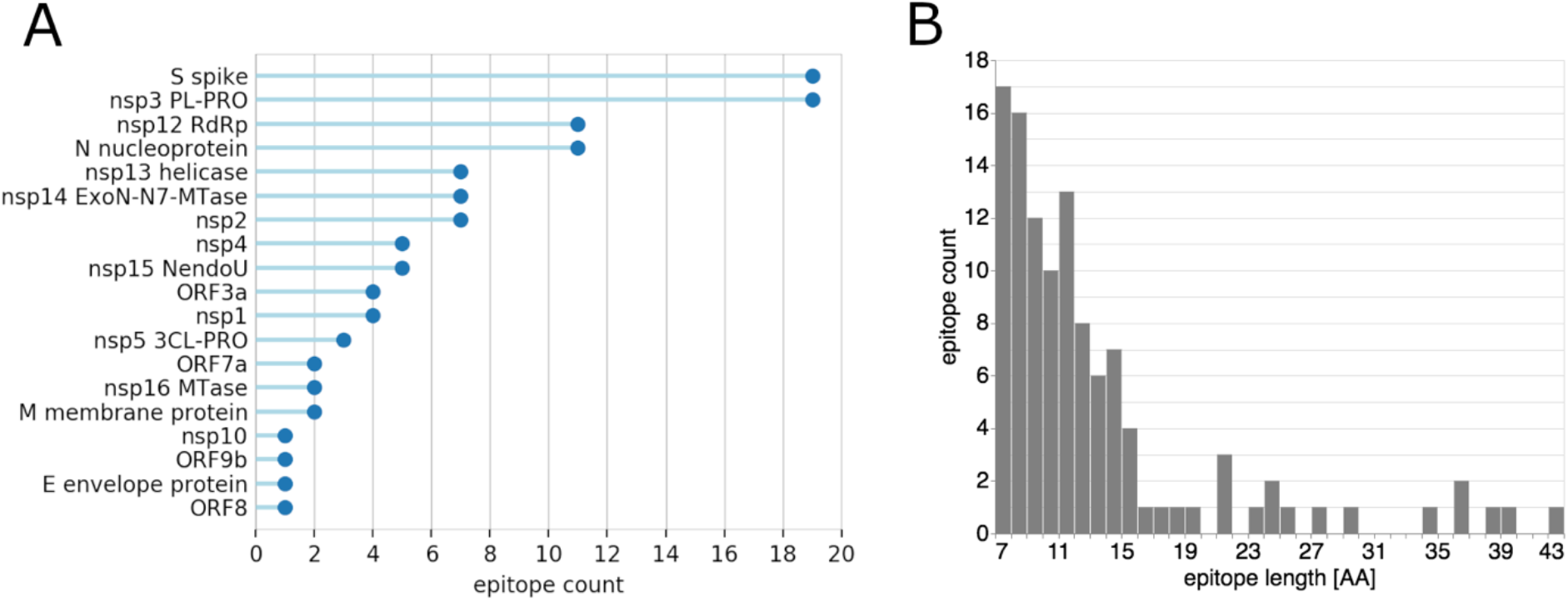
Linear epitope BepiPred2 predictions: **A**) summary count per protein with at least one predicted epitope; **B**) length distribution.

Second, we assessed the level of sequence conservation of each SARS-CoV-2 epitope compared to endemic human coronaviruses (HCoVs) strains HKU1, 229E, OC43 and NL63. For this purpose, a comprehensive database of endemic HCoVs protein sequences was built from 3 different sources: NCBI, UniProt and ViPR (**Table S2**). Prior to performing the comparison, each epitope was trimmed to 15 residues to avoid length bias. Trimmed epitopes were scanned using a sliding window against the endemic HCoV sequences. The average pairwise identity between the epitope and aligned residues was recorded at each position. Conservation was defined as the maximal value obtained after scanning was completed.

Conservation levels ranged between 40-100% with large local variations (**Figure S1 B**). Perhaps unsurprisingly, the most conserved epitopes were located in enzymes that perform essential functions in viral replication, such as the nsp12 RNA-dependent RNA polymerase, and are thus likely to be conserved across species (**Figure 2**). For proteins with less conserved epitopes overall, the average masked large local differences. In particular, the nucleoprotein contained epitopes in both the N-terminal RNA-binding and C-terminal dimerization domains, but only the C-terminus includes highly conserved, non-SARS-CoV-2-specific epitopes.

**Figure 2.**
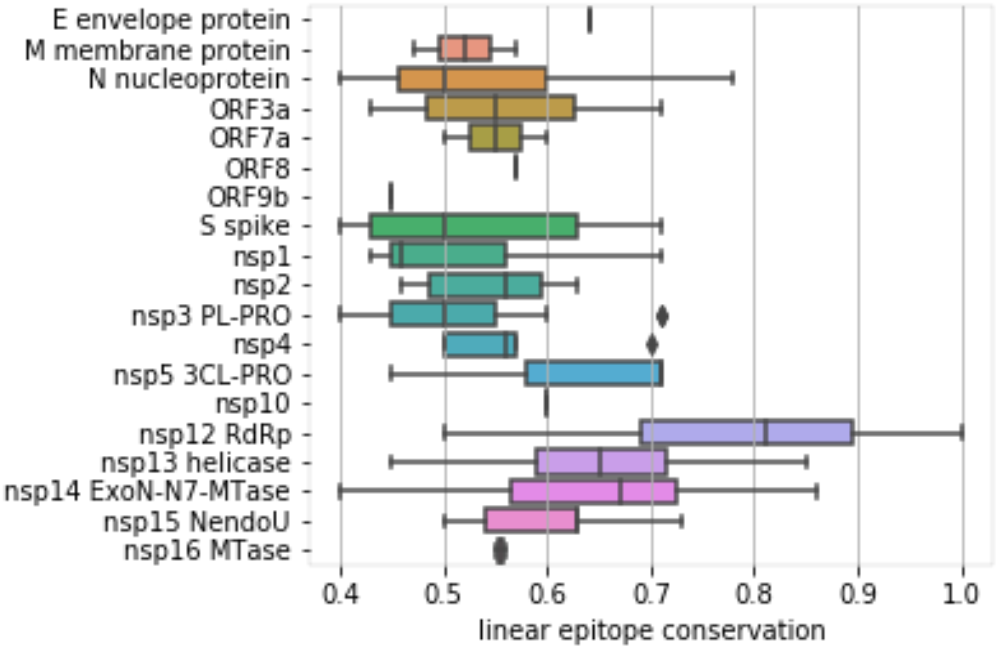
Distribution of conservation levels (1=100% identity) of SARS-CoV-2 linear epitopes compared to our comprehensive database of protein sequences from HCoV endemic strains. Epitopes located on essential enzymes such as the RdRp and helicase are the most conserved.

Third, conformational B-cell epitopes were predicted with DiscoTope2 on molecular structures of SARS-CoV-2 proteins^8^. The unprecedented speed at which experimental SARS-CoV-2 structures have been solved and deposited in the PDB allowed us to predict epitopes on 13 proteins (**Table S3**)^9–12^. Coverage was further extended to all proteins except E envelope protein, ORF9b and ORF14, by including high-quality homology models obtained with the Robetta platform and *ab-initio* models from the CASP-commons competition (**Table S3**)^13,14^. Since the Robetta models were based on PDB templates with high sequence similarity (e.g. SARS-CoV homologs with greater than 90% sequence identity), they are likely to have near atomic level accuracy in the range of less than a few angstroms root mean square deviation (rmsd). For example, the Robetta model for the closed state of the spike trimer is within 2 angstroms rmsd to the experimentally determined structure (6vxx) which was released after the models were generated. The CASP models are *de-novo* predictions and thus the prediction of conformational epitopes on these models are not directly based on experimental structure determination.

DiscoTope2 scores were plotted on the sequence to show the overlap between conformational and linear epitopes (**Figure S1 C**). Seven linear epitopes, one on nsp14, and six on spike, had no structural coverage because they matched gaps in the structures, which are indicative of highly flexible segments of the protein. Correlations between mobile segments and antigenicity are well known^15^. Provided they are not masked by another dominant epitope, we might therefore expect these epitopes to elicit a strong immune response even on the fully folded protein. One of these, 469-STEIYQAGST on the receptor-binding domain of the spike protein (spike RBD), could be of particular interest as conservation analysis indicates that it is SARS-CoV-2 specific.

To predict whether a linear epitope might be dominant, we assigned a ‘dominance score’ to regions where linear and conformational models both predicted an epitope. Six potentially dominant epitopes were thus predicted: nprot_5, nsp3_21 and nsp16_4 are likely to be specific with a conservation level below 0.6 (i.e specificity above 0.4); nsp2_4, nprot_10 and nprot_11 are likely non-specific (**Figure 3**). Interestingly, three of these epitopes are located on the nucleoprotein: the specific epitope is on the N-terminal RNA-binding domain, while the two nonspecific epitopes are on the C-terminal dimerization domain. This observation suggests that removal of C-terminal domain might be required to achieve specificity from endemic coronaviruses when assaying antibodies against the nucleoprotein.

**Figure 3.**
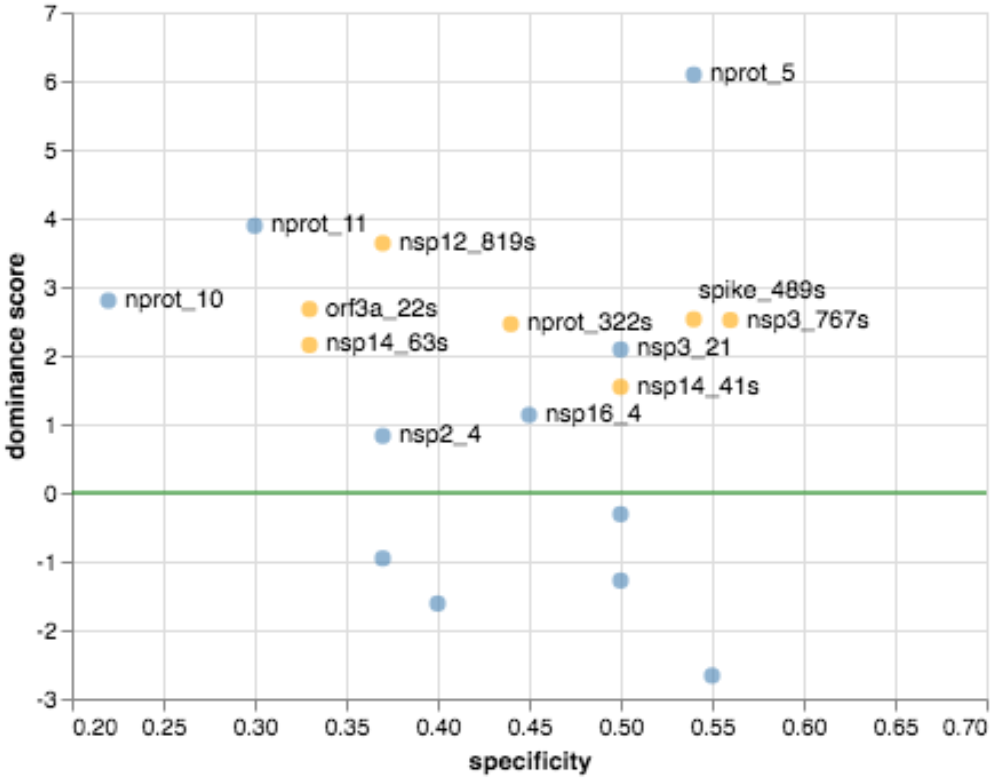
Dominance score *vs* specificity of predicted linear B-cell epitopes (blue) and structural epitopes (orange). Specificity is calculated as 1 - conservation. The dominance threshold (DiscoTope2 minimal score scaled to 0) is shown as a green line. Potentially dominant epitopes are labelled. Scores below −3 have been omitted for clarity.

Conformational epitopes were also predicted in regions where the linear epitope signal was below the threshold. Four of these structural epitopes were specific with a high dominance score, including nsp3_767s on the papain-like protease and spike_489s on the spike RBD. Lack of linear epitope signal could suggest that an immune response to these epitopes might only be achieved with the native protein or a structural mimic. However, it is also conceivable that the linear prediction tool simply failed to detect these epitopes.

Finally, we analyzed the SARS-CoV-2 genomes in the GISAID repository to locate variants that might affect our specificity predictions. A conservative approach was adopted to extract variants from an alignment of 6,796 unique GISAID sequences (downloaded on 04/20/2020), where each step sought to minimize erroneous variant calls due to sequencing errors. The degree of variability was further quantified by calculating the Shannon entropy at each position in the alignment. Variants were observed in all but 10 of the predicted linear epitopes (**Figure S1 D and E**). Unsurprisingly, no variants were observed in the nsp12_20 epitope that is 100% conserved in the endemic HCoVs. Shannon entropy identified eleven positions with conserved variants on 9 proteins. Sites with the highest Shannon Entropy values were nsp12 L323P and spike G614D, with a variant frequency of 40% in both cases. The spike variant is located in the 601-640 region that has been previously reported as 78% identical to a dominant B-cell epitope on SARS-CoV^6^. However, G614 is in a buried beta-sheet in the spike cryo-EM structure and no linear or structural epitope was predicted at that position. Two conserved variants were detected in predicted epitopes. The first is G251V on epitope orf3a_5 249-IDGSSGVVNP observed in 12% of ORF3a sequences; however, orf3a_5 is non-specific and is probably not a good diagnostic antigen candidate. The second is P586S on epitope nsp2_13 584-LQPLEQPTSEAVE observed in 5% of nsp2 sequences. Epitope nsp2_13 is specific; however, we found that the substitution had no effect on linear epitope prediction results or specificity. While it is straightforward to analyze the effect of single mutations, the exercise becomes more complex if multiple positions vary. To predict whether an epitope might be susceptible to sequence variations, we assigned a variant density score that reflects how many positions had at least one observed variant and compared this value to the overall variation frequency (**Figure 4**). The resulting plot reveals at a glance the epitopes with a conserved variant discussed above, orf3a_5 and nsp2_13. **Figure 4** also highlights three epitopes with infrequent variants (low SE100) but where 80% or more of positions are susceptible to variations, nprot_8_1, orf8_3, and nprot_322s. These epitopes might be too variable to consider for diagnostic use.

**Figure 4.**
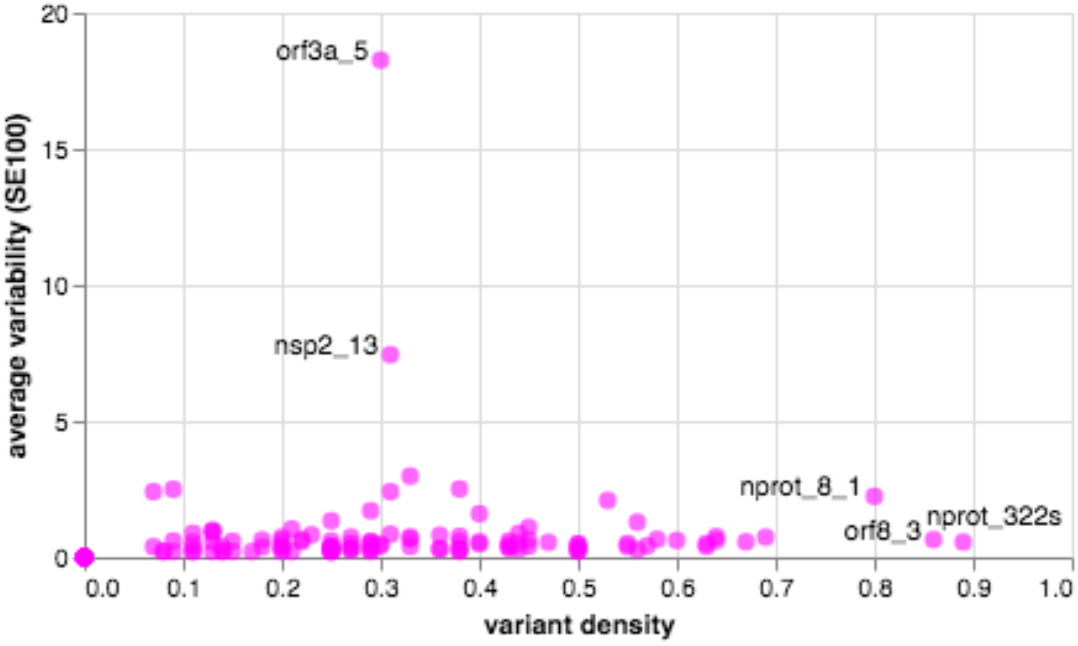
Amino acid variation in predicted epitopes, as average variability (SE100 = 100*Shannon entropy over epitope length) *vs* variant density. Epitopes with variants detected in over 80% of residues or variability above 5 are labelled. The high SE100 of orf3a_5 and nsp2_13 are due to single conserved variants.

### Epitope ranking

The summary plots provide a visual guide to evaluate the suitability of each protein for serological diagnostics, based on the raw predictions results displayed along the sequence. Favorable epitopes combine continuous BepiPred2 scores above the threshold, low conservation *vs* endemic HCoVs, continuous DiscoTope2 scores above the threshold, low variability and low density of variants. Our rationale for ranking individual epitopes based on derived data such as dominance and variant density scores is reflected in the scoring table (**Table S4**). Epitopes are classified into one of four broad categories: specific with high dominance score, specific linear, non-specific with high dominance score and non-specific linear. For each protein, we compiled the numbers of epitopes per category to predict whether the protein is likely or not to elicit an immune response and whether that response is likely or not to be specific (**Table 1**). As illustrated in the table, candidates with no predicted epitopes above the thresholds are marked as ‘no response’. Our top candidates marked as ‘specific’ contain only specific epitopes and at least one of them has a high dominance score. Three proteins match these criteria: the N nucleoprotein RNA-binding domain, the nsp3 papain-like viral protease domain and the nsp16 MTase. Next are the proteins that match the same criteria, but also contain non-specific linear epitopes: the S spike RBD and nsp3 full-length protein. We postulate that the immune response will likely be dominated by the specific epitopes with high dominance score as long as the protein is folded in native conformation. Altogether these 5 proteins would be our top candidates for serological diagnostics.

**Table 1.**
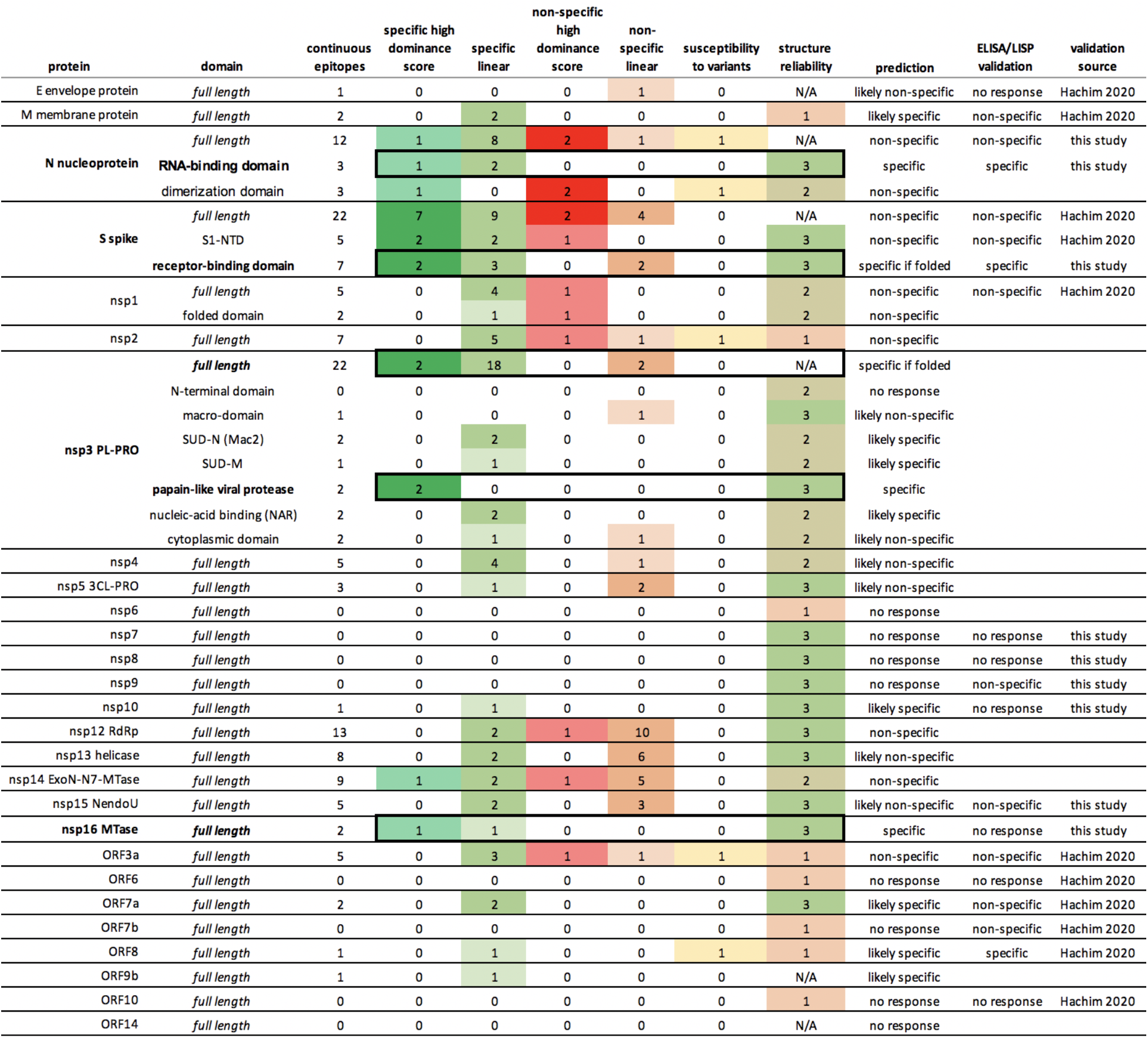
Summary predictions of SARS-CoV-2 diagnostic candidates, with the numbers of predicted epitopes in each category. Flexible segments are classified as having a high dominance score. Susceptibility to variants indicates the presence of epitopes with conserved mutations or a variant density above 80%. Structure reliability is set to 3 for experimental structures (very reliable), 2 for homology models and 1 for *de-novo* models (unreliable). Top five candidates are highlighted with a black box.

| protein | domain | continuous epitopes | specific high dominance score | specific linear | non-specific high dominance score | non-specific linear | susceptibility to variants | structure reliability | prediction | ELISA/LISP validation | validation source |
| --- | --- | --- | --- | --- | --- | --- | --- | --- | --- | --- | --- |
| E envelope protein | full length | 1 | 0 | 0 | 0 | 1 | 0 | N/A | likely non-specific | no response | Hachim 2020 |
| M membrane protein | full length | 2 | 0 | 2 | 0 | 0 | 0 | 1 | likely specific | non-specific | Hachim 2020 |
| N nucleoprotein | full length | 12 | 1 | 8 | 2 | 1 | 1 | N/A | non-specific | non-specific | this study |
|  | RNA-binding domain | 3 | 1 | 2 | 0 | 0 | 0 | 3 | specific | specific | this study |
|  | dimerization domain | 3 | 1 | 0 | 2 | 0 | 1 | 2 | non-specific |  |  |
| S spike | full length | 22 | 7 | 9 | 2 | 4 | 0 | N/A | non-specific | non-specific | Hachim 2020 |
|  | S1-NTD | 5 | 2 | 2 | 1 | 0 | 0 | 3 | non-specific | non-specific | Hachim 2020 |
|  | receptor-binding domain | 7 | 2 | 3 | 0 | 2 | 0 | 3 | specific if folded | specific | this study |
| nsp1 | full length | 5 | 0 | 4 | 1 | 0 | 0 | 2 | non-specific | non-specific | Hachim 2020 |
|  | folded domain | 2 | 0 | 1 | 1 | 0 | 0 | 2 | non-specific |  |  |
| nsp2 | full length | 7 | 0 | 5 | 1 | 1 | 1 | 1 | non-specific |  |  |
| nsp3 PL-PRO | full length | 22 | 2 | 18 | 0 | 2 | 0 | N/A | specific if folded |  |  |
|  | N-terminal domain | 0 | 0 | 0 | 0 | 0 | 0 | 2 | no response |  |  |
|  | macro-domain | 1 | 0 | 0 | 0 | 1 | 0 | 3 | likely non-specific |  |  |
|  | SUD-N (Mac2) | 2 | 0 | 2 | 0 | 0 | 0 | 2 | likely specific |  |  |
|  | SUD-M | 1 | 0 | 1 | 0 | 0 | 0 | 2 | likely specific |  |  |
|  | papain-like viral protease | 2 | 2 | 0 | 0 | 0 | 0 | 3 | specific |  |  |
|  | nucleic-acid binding (NAR) | 2 | 0 | 2 | 0 | 0 | 0 | 2 | likely specific |  |  |
|  | cytoplasmic domain | 2 | 0 | 1 | 0 | 1 | 0 | 2 | likely non-specific |  |  |
| nsp4 | full length | 5 | 0 | 4 | 0 | 1 | 0 | 2 | likely non-specific |  |  |
| nsp5 3CL-PRO | full length | 3 | 0 | 1 | 0 | 2 | 0 | 3 | likely non-specific |  |  |
| nsp6 | full length | 0 | 0 | 0 | 0 | 0 | 0 | 1 | no response |  |  |
| nsp7 | full length | 0 | 0 | 0 | 0 | 0 | 0 | 3 | no response | no response | this study |
| nsp8 | full length | 0 | 0 | 0 | 0 | 0 | 0 | 3 | no response | no response | this study |
| nsp9 | full length | 0 | 0 | 0 | 0 | 0 | 0 | 3 | no response | non-specific | this study |
| nsp10 | full length | 1 | 0 | 1 | 0 | 0 | 0 | 3 | likely specific | no response | this study |
| nsp12 RdRp | full length | 13 | 0 | 2 | 1 | 10 | 0 | 3 | non-specific |  |  |
| nsp13 helicase | full length | 8 | 0 | 2 | 0 | 6 | 0 | 3 | likely non-specific |  |  |
| nsp14 ExoN-N7-MTase | full length | 9 | 1 | 2 | 1 | 5 | 0 | 2 | non-specific |  |  |
| nsp15 NendoU | full length | 5 | 0 | 2 | 0 | 3 | 0 | 3 | likely non-specific | non-specific | this study |
| nsp16 MTase | full length | 2 | 1 | 1 | 0 | 0 | 0 | 3 | specific | no response | this study |
| ORF3a | full length | 5 | 0 | 3 | 1 | 1 | 1 | 1 | non-specific | non-specific | Hachim 2020 |
| ORF6 | full length | 0 | 0 | 0 | 0 | 0 | 0 | 1 | no response | no response | Hachim 2020 |
| ORF7a | full length | 2 | 0 | 2 | 0 | 0 | 0 | 3 | likely specific | non-specific | Hachim 2020 |
| ORF7b | full length | 0 | 0 | 0 | 0 | 0 | 0 | 1 | no response | non-specific | Hachim 2020 |
| ORF8 | full length | 1 | 0 | 1 | 0 | 0 | 1 | 1 | likely specific | specific | Hachim 2020 |
| ORF9b | full length | 1 | 0 | 1 | 0 | 0 | 0 | N/A | likely specific |  |  |
| ORF10 | full length | 0 | 0 | 0 | 0 | 0 | 0 | 1 | no response | no response | Hachim 2020 |
| ORF14 | full length | 0 | 0 | 0 | 0 | 0 | 0 | N/A | no response |  |  |

### Validation

We constructed a panel of 80 persons that had COVID-19 disease, all of whom reported having a positive nasopharyngeal swab for SARS-CoV-2 RNA. 71 of 80 persons in our cohort were convalescent with 21 days or more since symptom onset. Only two persons were likely acutely infected, one who had a blood draw the day of their SARS-CoV-2+ RNA test and another who had a blood draw 2 days after their SARS-CoV-2+ RNA test. The remaining 7 individuals were between 9 and 19 days of symptom resolution. We also gathered 106 serum and plasma “negative control” samples either collected before November 2019 (N=77) or demonstrated negative on the Abbott Architect SARS-CoV-2 IgG Assay (N=29).

From a subset of 6 COVID-19+ plasma and 5 “negative control” plasma, we compared our panel of recombinant SARS-CoV-2 proteins, including Spike-Receptor Binding Domain (S-RBD), Nucleoprotein (full length), nsp9, nsp10/nsp16 complex, nsp15, and nsp7/nsp8 complex (**Figure 5**). S-RBD and Nucleoprotein full length both gave significantly more ELISA signal with the COVID-19+ patients’ serum/plasma samples than the negative control samples. However, the signal-to-noise ratio of full-length nucleoprotein was only about 2, consistent with the predictions that the C-terminal portion of nucleoprotein has epitopes that are conserved with low pathogenicity coronaviruses. We then expressed the N-terminal, RNA-binding domain of the nucleoprotein (N-Nt), AA residues 47-173 and used this as an antigen for ELISA. The robust signal was retained in the N-Nt ELISA, and the signal-to-noise increased to 4, a 2-fold increase.

**Figure 5.**
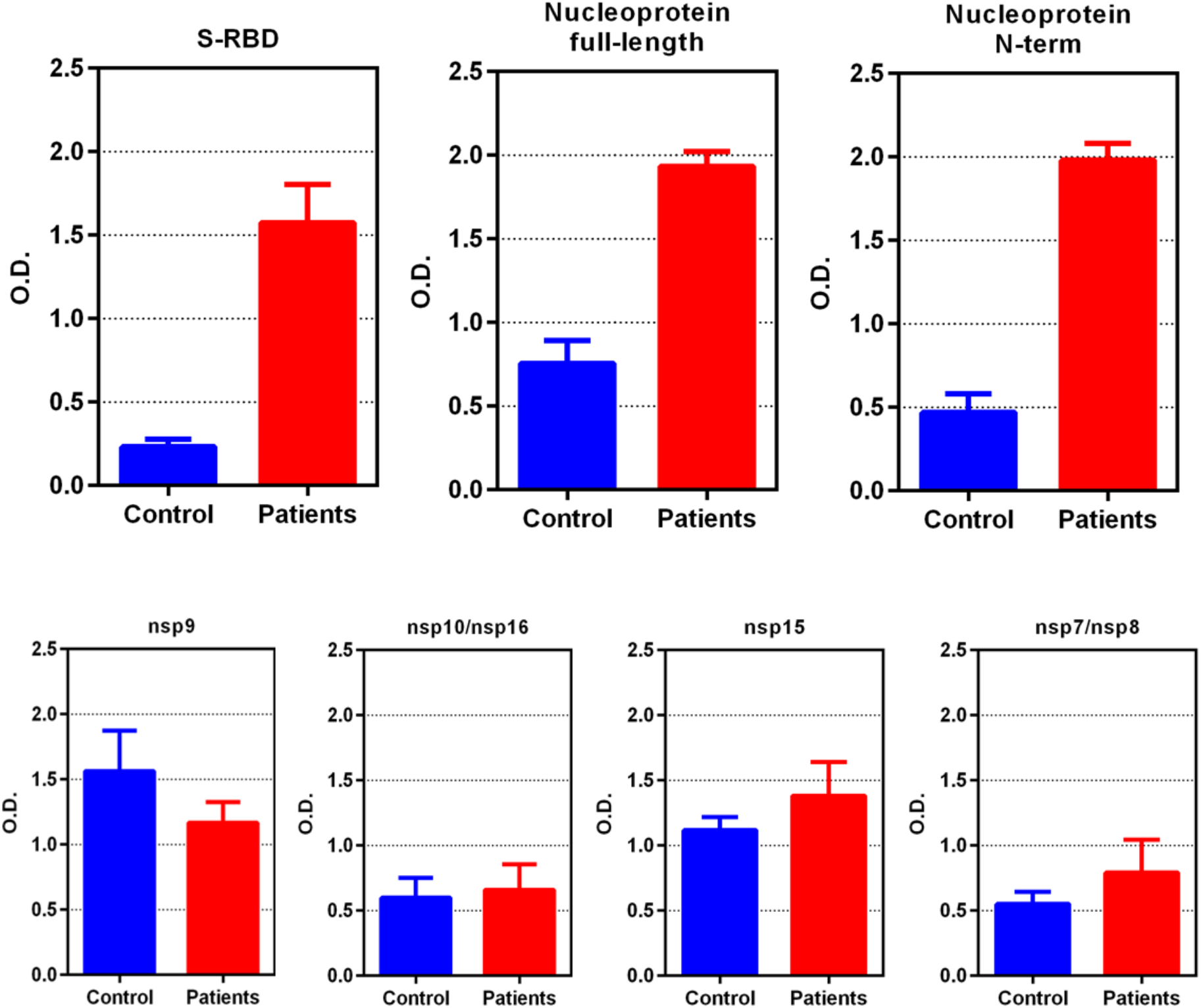
ELISA results (mean optical density at 492 [OD492], error bars are standard deviations of the mean) comparing plasma from 6 COVID-19 persons and 5 controls taken before November 2019 using anti-IgG conjugated to horseradish peroxidase (HRP). Performance of Spike-receptor binding domain (S-RBD) and Nucleoprotein in selective reaction with COVID-19 plasma IgG is far superior than nsp9, complexes of nsp7/8, complexes of nsp10/16, or nsp15. Note that the N-terminal portion of Nucleoprotein (N-Nt) retains the superior signal of full-length Nucleoprotein (Nuc) but improves the signal-to-noise ratio of the assay by nearly 2-fold, to 4.2 from 2.3, respectively. This implies that epitopes on the C-terminal portion of Nuc are non-specific for SARS-CoV-2, as predicted.

Testing 78 COVID-19+ sera/plasma, we found N-Nt gave a sensitivity of 87% (68 of 78 COVID-19+ positive); using 22 negative controls yielded 100% specificity. (**Table S5**). Testing S-RBD by ELISA yielded a sensitivity of 92% (72 of 78 COVID-19+ positive) and 100% specificity (0 of 22 negative controls tested positive). A few COVID-19+ persons were disconjugate in their seroreactivity, that is, were positive for one antigen in ELISA but negative for the other.

Combining the two antigens, S-RBD and N-Nt, to coat ELISA wells detected all of the positive sera in both assays including the disconjugate ones, but also revealed a couple of COVID-19+ plasma that were negative for both antigens into the positive range. Thus, this combined S-RBD and N-Nt ELISA was run on all 80 COVID-19+ sera/plasma and all 106 negative control sera/plasma yielding 94% sensitivity (75 of 80 COVID-19+ persons were positive) and 97.2% specificity (three of the 106 negative control plasma or sera tested were positive). The five negatives in the combined dual antigen assay were: one person with SARS-CoV-2 RNA positive test the same day (no records exist as to length of symptoms); a second person with COVID-19 symptom onset 17d prior, a single positive SARS-CoV-2 RNA test, then multiple negatives ensuing days; and three persons that had a positive SARS-CoV-2 RNA test (all adults >28d after symptom onset).

These results were combined with a recent report from an extensive serological study on SARS-CoV-2 proteins using LISP^4^ and compared to our in-silico predictions (**Table 1**). The experimental data appeared to validate our in-silico predictions for Nuc, N-Nt, spike S1, S-RBD, nsp1, nsp7, nsp8, nsp15, ORF3a, ORF6, ORF8 and ORF10. However, the LISP study and others have reported that Nuc is specific^2^. Our conservation analysis shows that the predicted dominant epitope nprot_10 on the C-terminal domain is matching at least 22 endemic HCoV strains from Seattle isolates, dated between 2015 and 2019, with 67% identity, which might explain some of the non-specific responses to full-length N in our samples (**Table S6**). The experimental data also contradicted our predictions for M, nsp16 and ORF7a. While M and ORF7a were not predicted to be good candidates, this was not the case for nsp16, which was in our top 5 selection. Although the ELISA was performed on the nsp10/nsp16 complex, the structure shows that the putative dominant epitope on nsp16 is in a loop away from the interface with nsp10 and therefore the lack of response cannot be attributed to masking due to complex formation. The remaining validation on E, nsp9, nsp10 and ORF7b was deemed inconclusive due to weak signal. Full length spike has been recently reported as specific in other studies^3^ and further validation is needed.

B-cell antigenicity is influenced by the antigen abundance during infection and the way it is displayed to the immune system. Our prediction method does not address the former and is only a crude approximation of the latter. For example, the conservation of discontinuous epitopes was not considered in this study as it involves the comparison of surface areas, which might be disrupted by local insertions. Current in-silico predictions suffer from sensitivity issues and this will only improve with larger training sets, i.e. more experimental structures of antigen-antibody complexes^8^. Nonetheless, our combined predictions accurately ranked the spike RBD and N-term nucleoprotein as top choice candidates for specific SARS-CoV-2 diagnostics, which was subsequently confirmed by ELISA serological tests. This prediction method can speed the design of serodiagnostics for future pandemics, as demonstrated in this test case for SARS-CoV-2/COVID-19 diagnostics. It can in principle incorporate results from any epitope prediction tool, as long as results are reported as one discrete value per residue. Raw data for estimating the effects of variants can alternatively be sourced directly from public databases, such as NextStrain^16^, ViPR^17^ and the CNCB^18^. However, due to data restrictions at GISAID, sequence alignments cannot be retrieved from those resources; we chose to perform our own variant analysis to be able to inspect protein alignments and retrieve strain information from individual sequences.

The ELISA described is a low-cost method to sensitively and specifically detect antibody to SARS-CoV-2. Both antigens are high expressors, though one is made in *E. coli* (N-Nt) and one in mammalian cells (S-RBD). We estimate the cost to be only pennies per test. In comparison, the total cost of commercial tests to the patient ranges from $40 to $140 per test, including $10/test for the reagents alone. We are in the midst of shipping this test to eight low-to-middle income countries as a cost-effective test to monitor their infection rates and for research purposes. Further validation to address specificity is required prior to use for patient management, especially in low prevalence populations. However, the low frequency of reactivity in 106 individuals suggests that antibodies to more frequent viruses such as influenza, RSV, CMV, HSV and others are not cross-reactive with these antigens.

### Reagent availability

Samples of N-Nt proteins have been deposited in the BEI Resources repository and can be acquired from https://www.beiresources.org/. The amount of N-Nt in one tube (1mg) would support the coating of 10,000 wells (individual assays, if not in duplicate or triplicate) for ELISA testing. Deposition of S-RBD is pending.

### Data and code availability

The file containing the raw data and the associated code to produce all the charts in Figure S1 are available on https://github.com/ssgcid/sc2_epitopes.

## Materials and methods

### Linear epitope analysis

Unless otherwise specified, the reference genome throughout this study is from the severe acute respiratory syndrome coronavirus 2 isolate Wuhan-Hu-1 (GenBank: MN908947.3). The corresponding protein sequences were selected as follows: polyprotein ORF1ab (UniProt accession: P0DTD1), Spike (P0DTC2), ORF3a (P0DTC3), E envelope protein (P0DTC4), M membrane protein (P0DTC5), ORF6 (P0DTC6), ORF7a (P0DTC7), ORF7b (P0DTD8), ORF8 (P0DTC8), N nucleoprotein (P0DTC9), ORF9b (P0DTD2), ORF10 (A0A663DJA2) and ORF14 (P0DTD3). The 15 mature products of polyprotein cleavage, nsp1, nsp2, nsp3 PL-PRO, nsp4, nsp5 3CL-PRO, nsp6, nsp7, nsp8, nsp9, nsp10, nsp12 RdRp, nsp13 helicase, nsp14 ExoN-N7-MTase, nsp15 NendoU and nsp16 MTase, collectively referred to as non-structural proteins (NSPs), were obtained from the chain features annotation of polyprotein ORF1ab on ViralZone^19^.

The comprehensive database of endemic human coronavirus (HCoV) protein sequences was built from 3 sources: NCBI, UniProtKB and ViPR. The UniProtKB and NCBI Identical Protein Groups databases were searched by the NCBI taxonomy identifiers for human coronaviruses strains HKU1, 229E, OC43 and NL63. Non-human strains sequences from taxonomy subnodes of strain 229E were excluded. Sequence variants manually annotated in the UniProtKB Swiss-Prot entries were extracted with VARSPLIC^20^. ViPR sequences were obtained by searching the taxonomy tree on the Coronaviridae Gene/Protein search form for strains HKU1, 229E and NL63, and by performing a search by keyword for [“ OC43” or “ HCoV-OC43”], followed by parsing of the Organism and Strain fields from the fasta headers and grouping by values. Nonhuman coronavirus sequences and fasta records with no protein sequences were discarded. Although NCBI, UniProtKB and ViPR datasets overlap, we found that each contained unique sequences: 1 for NCBI, 12 for ViPR and 51 for UniProtKB. These unique sequences could potentially influence results since a single variant on a 7-residue epitope has a 14% impact on the conservation score. The combined dataset of 15,188 sequences was reduced to 1,740 after redundancy elimination and clustering with CD-HIT to remove 100% overlapping fragments^21^. Polyprotein sequences were identified by blastp similarity searches against the reference and NSPs extracted to avoid false predictions across boundaries, yielding 1,138 unique NSP sequences. These were added back to the non-polyprotein dataset. After removal of 27 sequences with known artificial mutations (originating from the PDB), the final HCoV reference database contained 2,234 protein sequences.

Linear B-cell epitopes were predicted for the 12 non-polyproteins and 15 NSPs of the reference SARS-CoV-2 isolate with BepiPred2.0, using a minimum length of 7 amino acid residues and minimum score of 0.55 (80% specificity/30% sensitivity)^5^. To reduce the chance of introducing bias towards lower conservation score due to length, long epitopes were trimmed or split to 15 residues prior to scanning for conservation against the endemic HCoV database, by increasing the local BepiPred2 threshold until the stretch of continuous high scores was at most 15 amino acid long. Epitope conservation was estimated by aligning the epitope against each sequence in the endemic HCoV database using a sliding window and calculating the pairwise identity at each position, averaged over the length of the epitope. Epitope conservation was defined as the maximal pairwise identity value obtained after scanning was completed.

### Conformational epitope analysis

Coordinates for experimental structures of SARS-CoV-2 proteins were downloaded from the PDB. In case multiple structures were available, the apo structures with highest sequence coverage and highest resolution were selected. To ensure the highest possible coverage of the SARS-CoV-2 structome, a repository of homology models was built at the onset using Robetta (https://www.ssgcid.org/cttdb/molecularmodel_list/?target_icontains=BewuA). Where no PDB structure was available, high confidence (overall confidence score > 0.5, local coordinate error < 3Å) Robetta models were used instead. In case no template was available for homology modelling, de-novo models were obtained from CASP-commons (https://predictioncenter.org/caspcommons/). Top scoring models as ranked by ProQ3D EMA^22^ were selected that also had the highest local score at epitope locations.

Conformational B-cell epitopes were predicted with DiscoTope2 with a significance threshold of −2.5 (80% specificity/40% sensitivity)^23^. Scores were scaled at zero for plotting. The dominance score for linear epitopes was defined as the DiscoTope2 score averaged over the length of the linear epitope. Epitopes that were discontinuous due to gaps in the structure were excluded. Conformational epitopes with no corresponding linear epitope were inspected for regions with continuous signal. Dominance and conservation were calculated over these regions if they spanned at least 6 residues. Regions longer than 15 residues with diffuse boundaries were excluded.

### SARS-CoV-2 variant analysis

A total of 7,948 high coverage sequences from human isolates were downloaded from GISAID on 04/20/2020, representing 6,797 unique DNA sequences. These were annotated as ‘sequences with <1% Ns and <0.05% unique amino acid mutations (not seen in other sequences in the database) and no insertion/deletion unless verified by the submitter’. The unique sequences were aligned to the reference genome with MAFFT experimental version 7.463 using options ‘--auto --addfragments’^24,25^. The misaligned 3.2 kb fragment EPI_ISL_426413 was identified as Hepatitis B virus isolates JRC-HB01 by MEGABLAST against the NCBI nr database and was removed from the alignment. Further, a total of 40 inserts in the alignment that introduced gaps in the reference sequence were deleted as likely sequencing errors. The 11 largest inserts ranged from 99 to 155 bases, and the next largest were 3 base long. The genome submitter confirmed that those large inserts were likely assembly artefacts that will be corrected. Of the smaller inserts, a three base ‘TTT’ in-frame insertion was observed at nsp6 position 298 in 17 sequences, and is presumably real, although it is in a repetitive region of 8 consecutive Ts and therefore at the limit of accurate PCR amplification of mono-thymidine repeats^26^. Gaps in aligned sequences were likewise treated as likely sequencing errors and replaced with the unknown base to keep translation in frame with respect to the reference sequence, taking into account the ribosomal frameshift in the nsp12 coding sequence. This process resulted in just 95 remaining premature terminations in over 160K proteins; 43 of these were found consistently after amino acid 125 of nsp10 and likely represent true truncations. The others were distributed across the sequences of 11 protein families and are likely erroneous. The 27 protein alignments thus obtained, one for each of the SARS-CoV-2 proteins selected for this study, were transformed into a n by m matrix, where n is the length of the reference protein and m the length of the alignment, and processed columnwise. At each position in the alignment, the variant count was obtained by counting unique amino acids that differed from the reference; variability was quantified using the Shannon entropy, calculated as 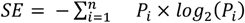 where n is the number of amino-acid types and Pi is the fraction of amino acid type i at that position^27,28^. Codes for unknown amino acid (X) and translation termination (*) were ignored in all calculations.

### Epitope scoring table

Supplementary Materials (**Table S4**). Epitopes with the highest dominance score and highest specificity were ranked at the top: first the epitopes defined by both the linear and conformational predictions; second, the conformational epitopes where the linear prediction was below the threshold; third, the specific dominant epitope with high variant density. This group was followed by epitopes located on flexible segments, ranked by specificity. The remaining linear epitopes were ranked by specificity, average BepiPred2 score and variant density. At the bottom of the list were the least specific epitopes on flexible segments, followed by those with the highest dominance score.

### Protein expression and purification

pET28 vector constructs encoding full length SARS-CoV-2 Nucleoprotein, nsp7, nsp8, nsp9, nsp10, nsp15 and nsp16 were ordered from Genscript. The nucleotide sequences were codon optimized for *E.coli* expression and in frame with an N terminus 6XHistidine metal affinity tag with a TEV protease cleavage site. Additionally, a pMCSG53 vector construct encoding the N terminus of SARS-CoV-2 Nucleoprotein (47-173), with an N terminal 6X Histidine metal affinity tag and TEV protease cleavage site was gifted from the Center for Structural Genomics of Infectious Diseases (CSGID).

Plasmids were transformed into BL21 (DE3) Rosetta Oxford chemically competent *E.coli* cells, and selected colonies were re-streaked on LB agar with appropriate antibiotics.

For protein production, starter cultures of PA-0.5G [50 mM Na_2_HPO_4_ 50 mM KH_2_PO_4_ 25 mM (NH_4_)2SO_4_ 2 mM MgSO_4_ 0.2x trace metals 0.5% glucose 200 μg/mL 18 amino acids] noninducing media with appropriate antibiotics were grown for 18 h at 25 °C. Antibiotics were added to 2 L bottles of sterile ZYP-5052 [1% tryptone 0.5% yeast extract 50 mM Na_2_HPO_4_ 50 mM KH_2_PO_4_ 25 mM (NH_4_)2SO_4_ 2 mM MgSO_4_ 0.2x trace metals 0.5% glycerol 0.5% glucose 0.2% lactose] autoinduction media, and the bottles were inoculated with overnight cultures^29^.

Inoculated bottles were then placed into a LEX bioreactor and cultures grown for 72 h at 20°C. To harvest, the medium was centrifuged at 6000 RCF for 30 min at 4°C. Cell paste was frozen and stored at −80 °C prior to purification.

Nsp7, nsp8, nsp9, nsp10, nsp16, nsp15, full-length nucleoprotein and the N-terminal truncation representing the RNA-binding domain (amino acids [AA] 47-173) of nucleoprotein (N-Nt) were all produced in *E. coli* and complexes assembled by co-lysis of recombinant proteins produced in individual *E. coli*. For purification, cells were resuspended in lysis buffer [25 mM HEPES (pH 7.0) 500 mM NaCl 5% glycerol 0.5% CHAPS 30 mM imidazole 10 mM MgCl_2_ 1 mM TCEP 1 mM AEBSF 0.025% NaN3] and lysed by sonication. The cell lysate was then cleared by centrifugation at 26000 g for 45 min at 4°C. Each recombinant protein was purified from the cell lysate by immobilized metal affinity chromatography. A Ni-NTA column (GE HisTrap FF) was equilibrated in lysis buffer to which the lysate was added. The column was washed with 150 mL of wash buffer supplemented with [25 mM HEPES (pH 7.0) 500 mM NaCl 5% glycerol 30 mM imidazole 1 mM TCEP 0.025% NaN3] and the protein was eluted from the column by the addition of wash buffer supplemented with 350 mM imidazole. In each case, the protein was further purified by size-exclusion chromatography on a GE HiLoad 26/600 Superdex 75 PG or a Superdex 200 PG column previously equilibrated with size-exclusion buffer [25 mM HEPES (pH 7.0) 500 mM NaCl 5% glycerol 2 mM DTT 0.025% NaN3]. Fractions containing the final protein, in a pure state, were pooled and concentrated to 5 mg/mL before being snap frozen in liquid nitrogen and stored at −80°C.

The prefusion-stabilized trimeric Spike (S-RBD) construct was produced at the Institute for Protein Design, Seattle, WA. The expression plasmid was received from David Veesler, UW Biochemistry, Seattle, WA, with discovery and characterization information as previously described^11^. The spike protein was expressed using transient transfection in Thermo Expi HEK293 mammalian cells per IPD protocol, harvested after 3 days incubation at 33°C, and clarified using PDAD (Sigma-Aldrich #409014). The clarified harvest fluid (CHF) was purified using TaKaRa Talon immobilized metal affinity chromatography (TaKaRaBio 635504) and buffer exchanged into Tris Buffered Saline at 2-8°C. The protein was characterized using negative stain Electron Microscopy, SDS-PAGE, and UV-Vis.

### Serological assays (ELISA validation)

Recombinant proteins refer to SARS-CoV-2 nsp7/8, nsp9, nsp10/nsp16, Spike-RBD, Nucleoprotein full length, N-Nt, and a mix of S-RBD and N-Nt. The recombinant proteins were diluted to 2 μg/mL into sterile 1x PBS and then used to coat ELISA plates (Immulon 4HBX Flat Bottom Microtiter Plates 96-wells, 50 plates/case, ThermoScientific, Rochester, NY), using 100μL of 2 μg/mL into each well, covered, and incubated at 4°C at least overnight (up to seven days). For the combined ELISA, we coated with both S-RBD at 1 μg/mL and N-Nt at 1 μg/mL in 100 μL PBS in each ELISA well. After incubation at 4°C at least overnight, the plates were washed with a plate washer 2x with 300μL per wash (wash solution for this and all washes: PBS with 0.1% Tween20). Then we added 200μL of 2% BSA (20 mg/mL) in PBS to each well to quench binding, incubating at least 1 hr at room temp with shaking. We then washed with a plate washer 2x at 300 μL per wash. We heat treated all plasma or sera at 56°C for 60 min. Tubes were cooled on ice after heat treatment and spun at 10,000 RPM in a 4°C microfuge for 10 min, saving the supernatant. The supernatant was then diluted to working concentration (1:100) in PBS-0.1% Tween20-2% BSA and spun again at top speed in a 4°C microfuge (13,000 RPM or 17,000 x g) for 30 min, and the supernatant was recovered for the next step. We then added 50μL per well of 1:100 diluted plasma or sera, diluted in PBS-0.1% Tween20-2% BSA and then incubated for 90 min at room temp, while shaking. We then washed the plates with a plate washer 3x 300μL per wash. We then added 100 μL per well of anti-IgG-HRP (Rockland anti-human-IgG(H+L) goat antibody horseradish peroxidase (HRP) conjugated, Cat# 609-1302) (diluted in same buffer as the sera/plasma), and incubated 1 hr at room temp while shaking. After that incubation, we washed with a plate washer 3x at 300μL per wash. We then added 100μL of developing solution (SigmaFAST OPD tablets, Sigma, St. Louis, MO, prepared as per manufacturer), allowed 30min in the dark at room temperature, and added 25uL stop solution (3N HCl). The plates were then covered and read on plate reader at OD 492nM. Positives were defined as those with OD492 that are greater than the (mean + 3 times standard deviation of mean) of the negative controls. The ELISA was run each time with 10 to 20 control plasma/sera for this calculation.

The samples from those persons that tested positive for SARS-CoV-2 were derived from plasma or sera from adults (>18 yo) of both genders, provided without personal identifiers, from Seattle Children’s SARS2 Recovered Cohort (N=16) and a convalescent cohort screened for eligibility for plasma donation (N=57). In addition, seven specimens were residual clinical samples obtained from inpatients at the University of Washington. Forty-two negative control plasma and serum samples were from banked samples collected from healthy male and female adult (>21 yo) volunteers who had previously participated in PK studies conducted earlier than November 2019 by Dr. Isoherranen’s group. A second group of negative control specimens included thirty sera samples from adults participating in University of Washington Virology Research Clinic protocols studying zoster vaccination earlier than November 2019. A third group of negative controls were 29 serum specimens from the Department of Laboratory Medicine that tested negative for COVID-19 in the Abbott Architect SARS-CoV-2 IgG Assay^30^. All negative control samples were obtained earlier than November 2019. All serum and plasma samples were collected under the University of Washington or Seattle Children’s IRB approval.

## Acknowledgements

We gratefully acknowledge the authors, originating and submitting laboratories of the sequences from GISAID’s EpiFlu™ Database on which this research is based. The list is detailed in Suppl. **Table S7**. We thank Christopher L. McClurkan and Victoria L. Campbell for assistance with specimen processing, Ryan Choi for help with ELISA graphs, all of SSGCID for excellent support, as well as Elisabeth Gasteiger and Richard Scheuermann for helpful discussions. We thank the University of Washington, Department of Laboratory Medicine, Northwest Biospecimens Repository for providing serum and plasma samples. We also thank The Audacious Project at the Institute for Protein Design for supporting the Robetta infrastructure and the CASP Commons competitors and organizers for making the *de-novo* models openly accessible.

## Supplemental information

**Suppl. Figure S1.**
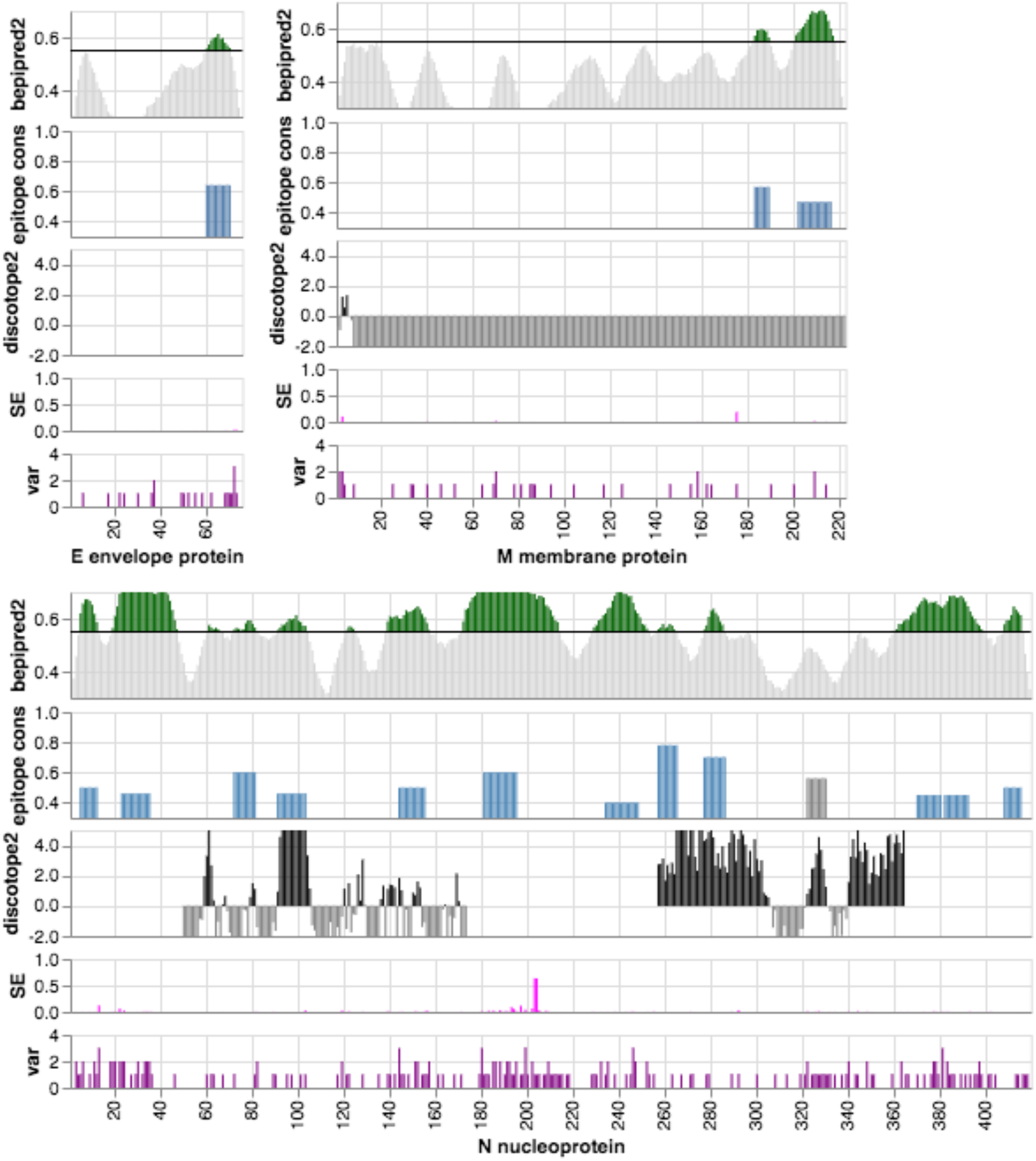
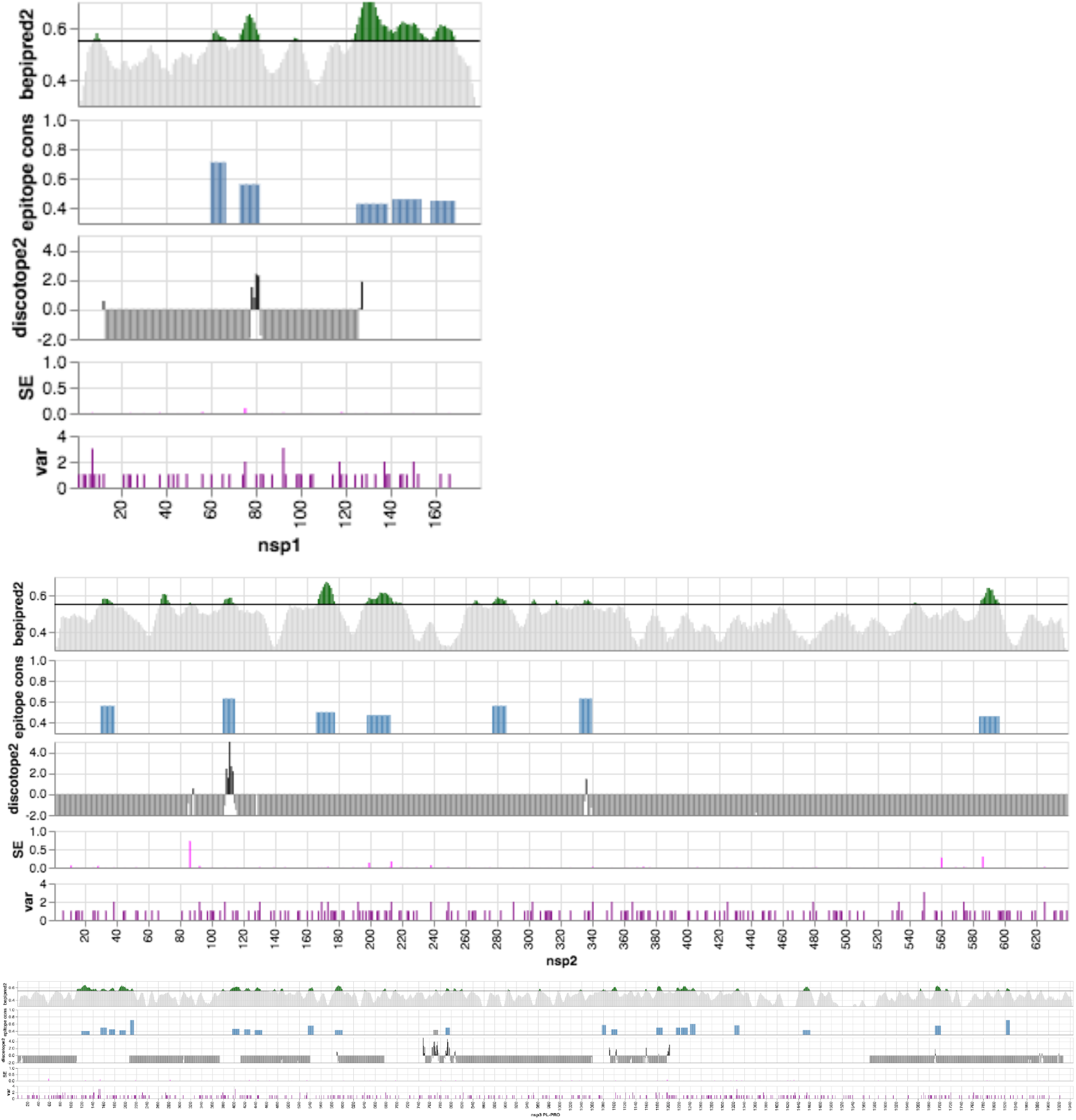
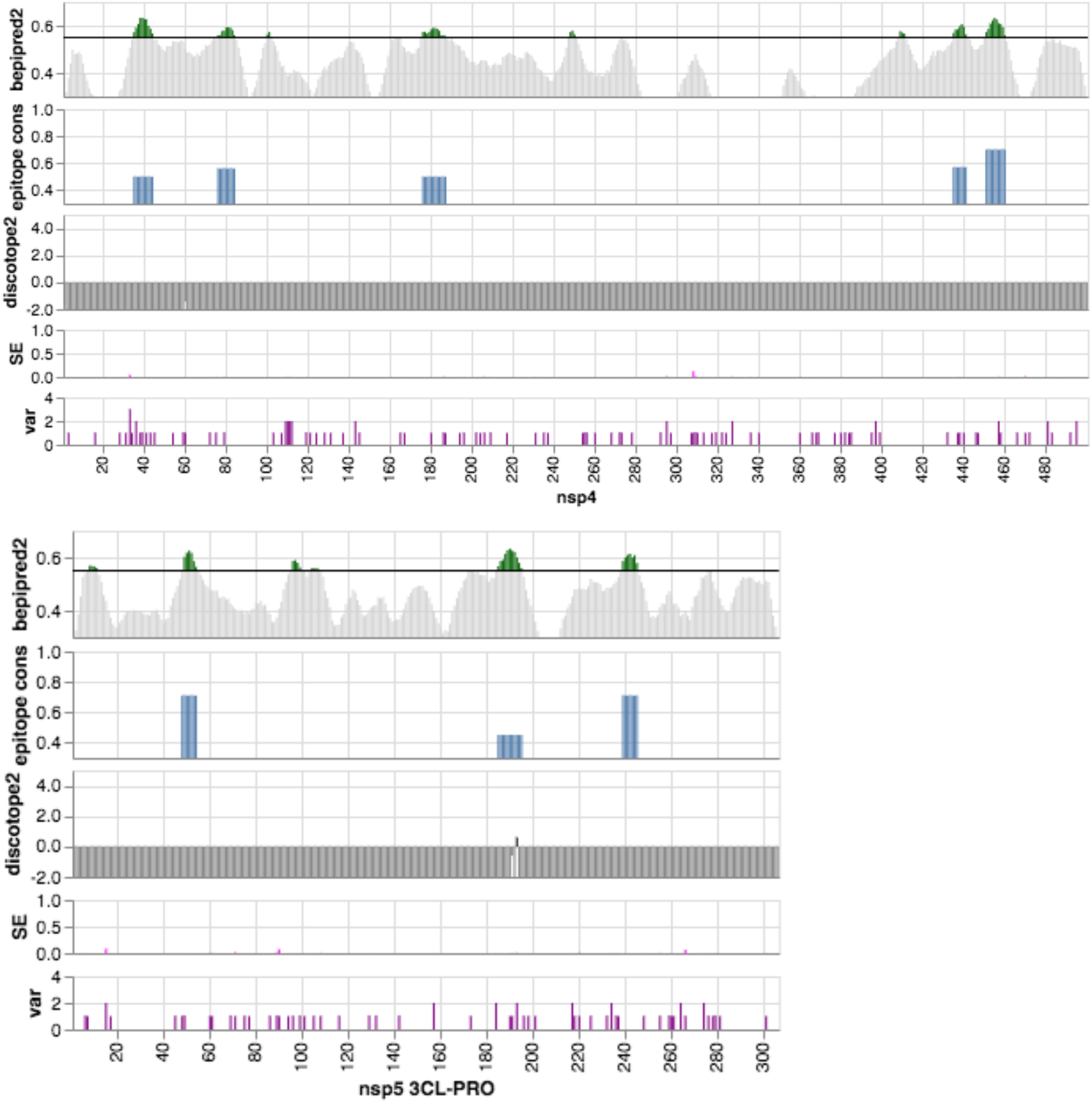
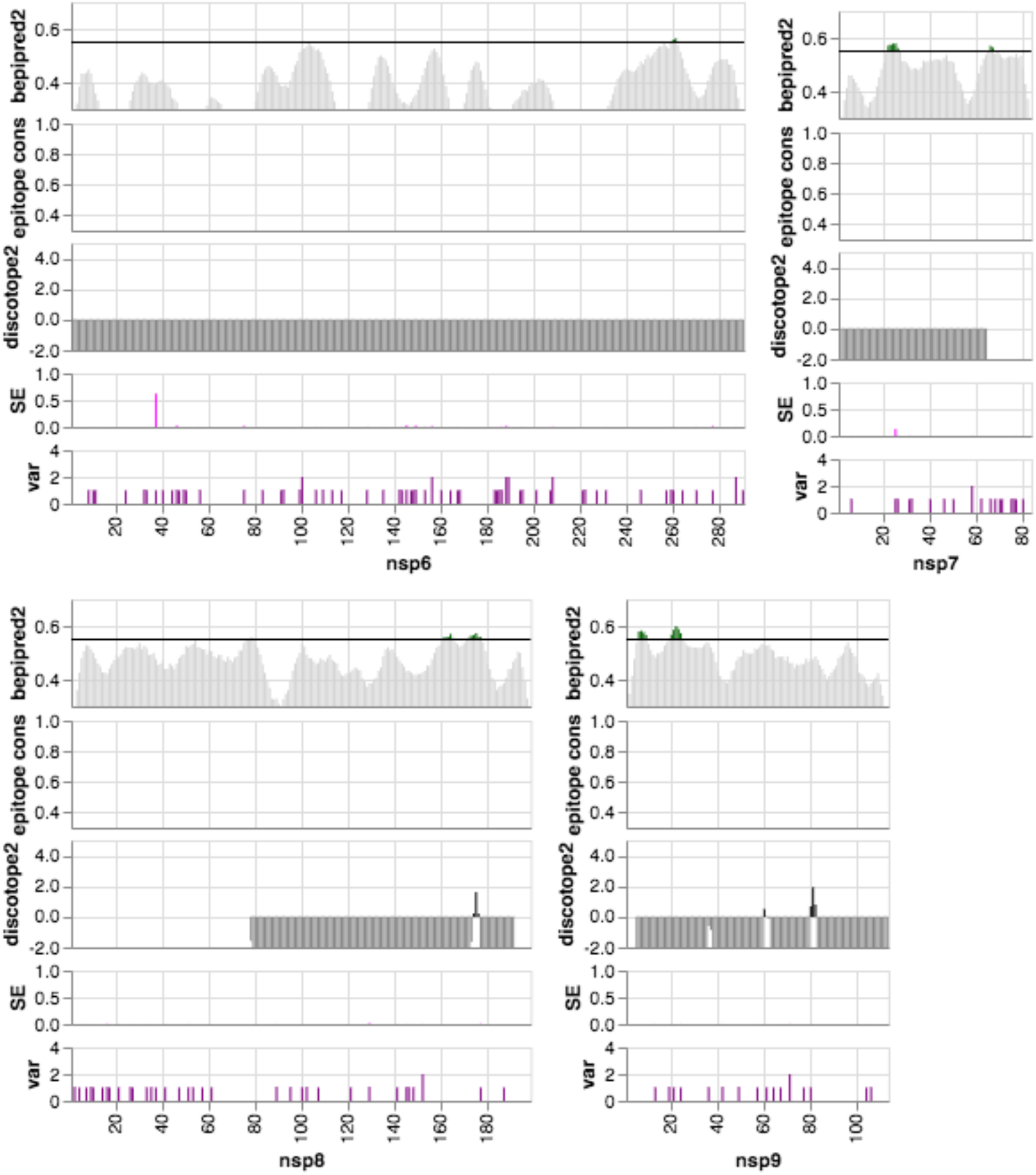
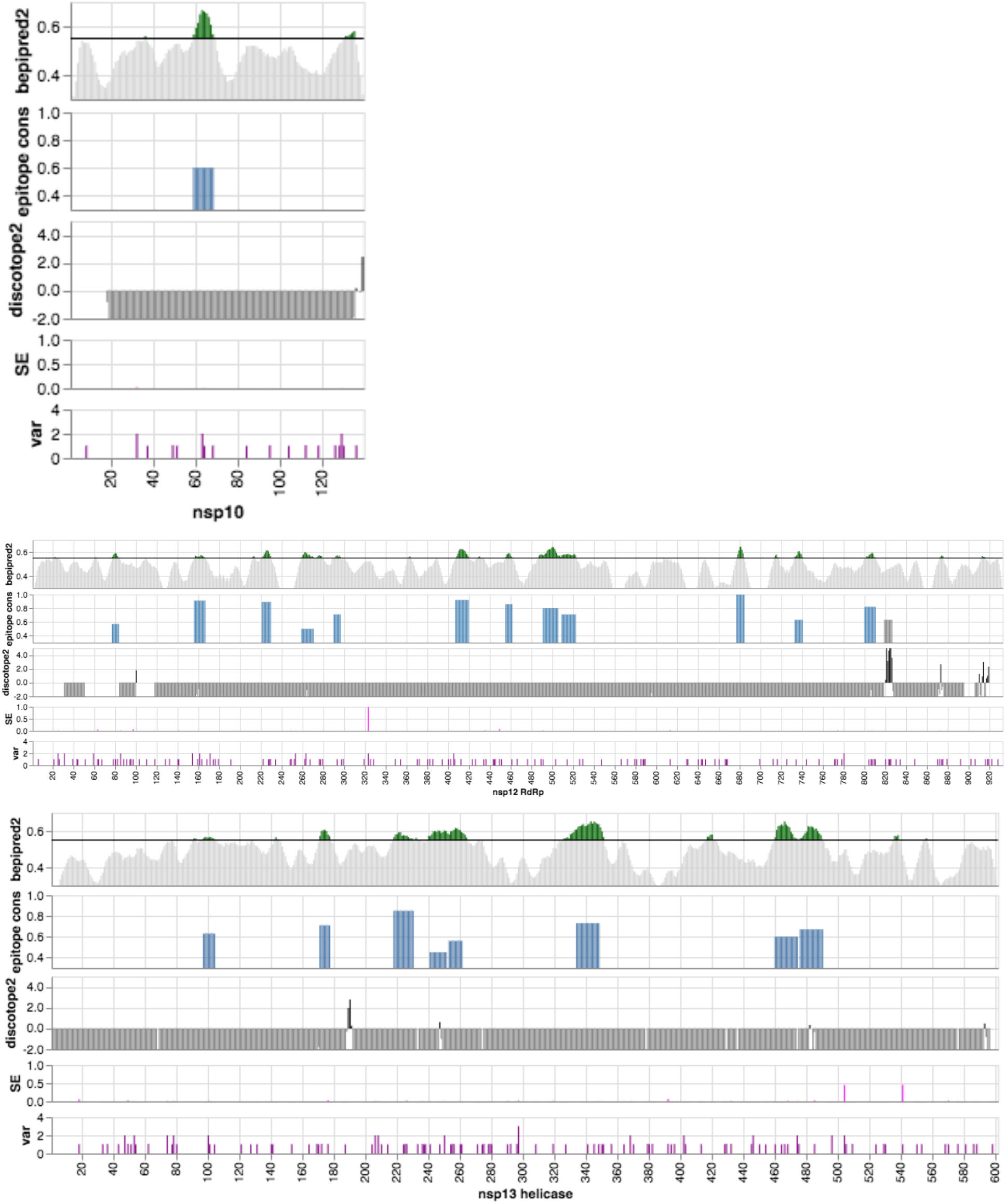
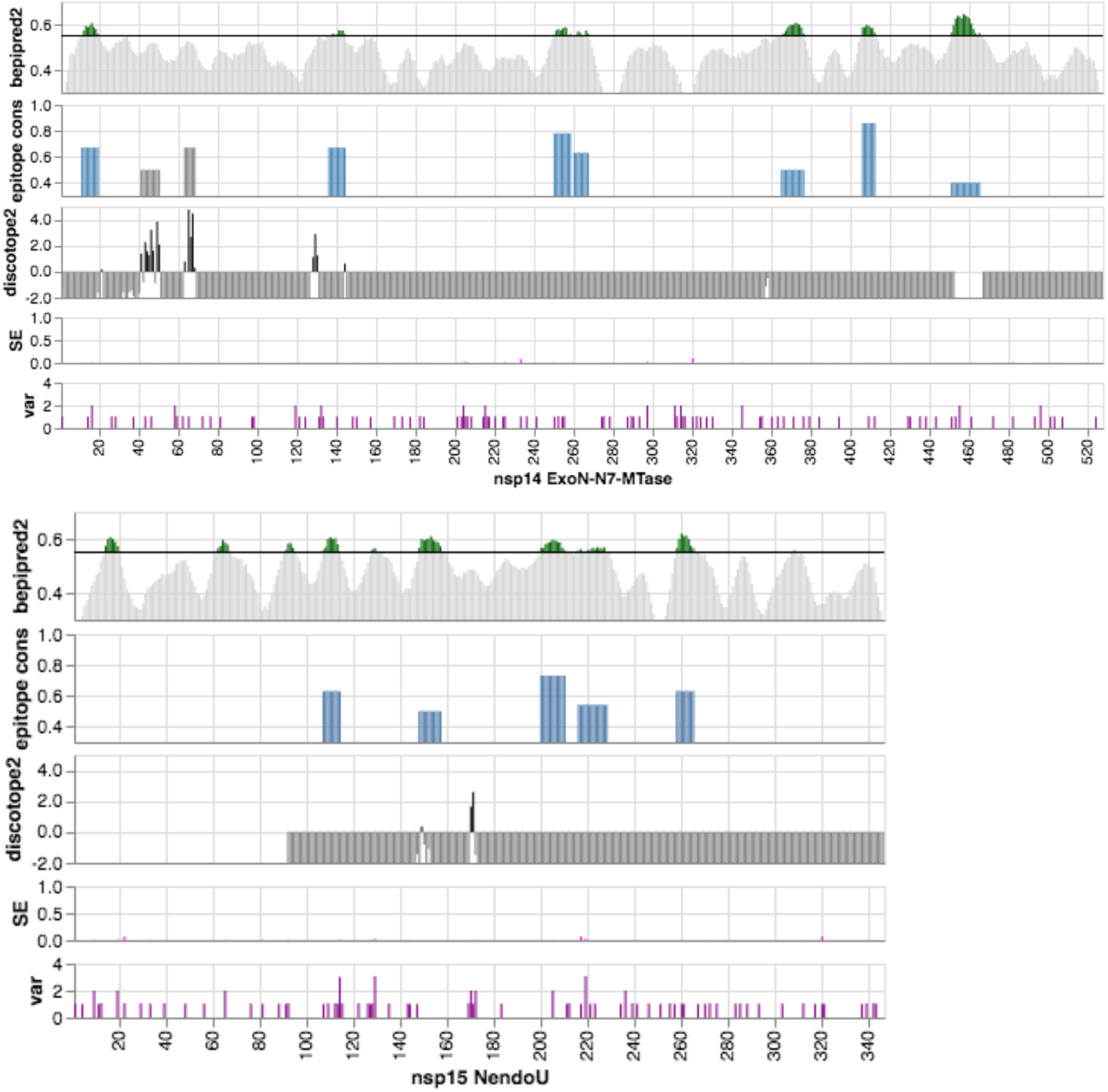
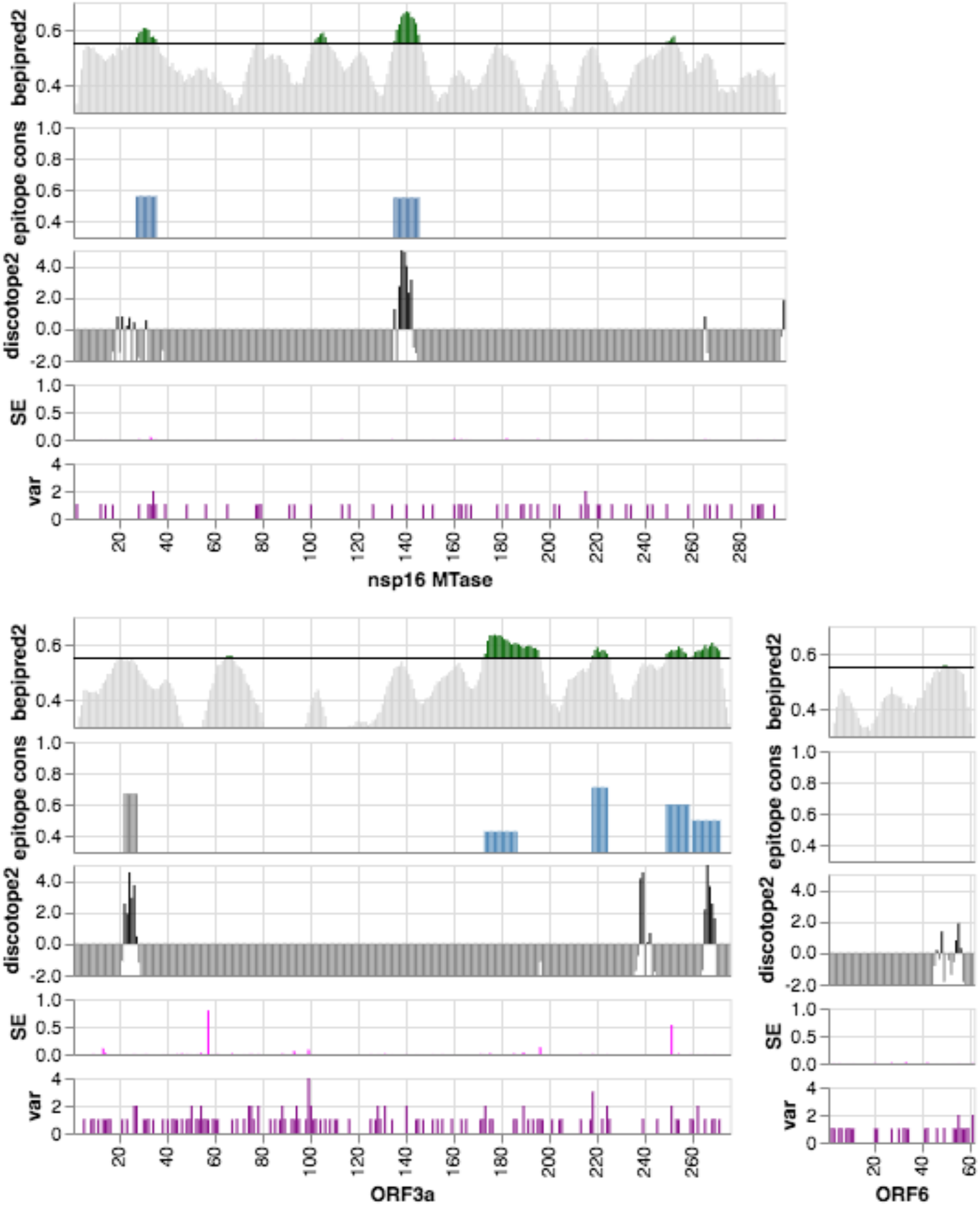
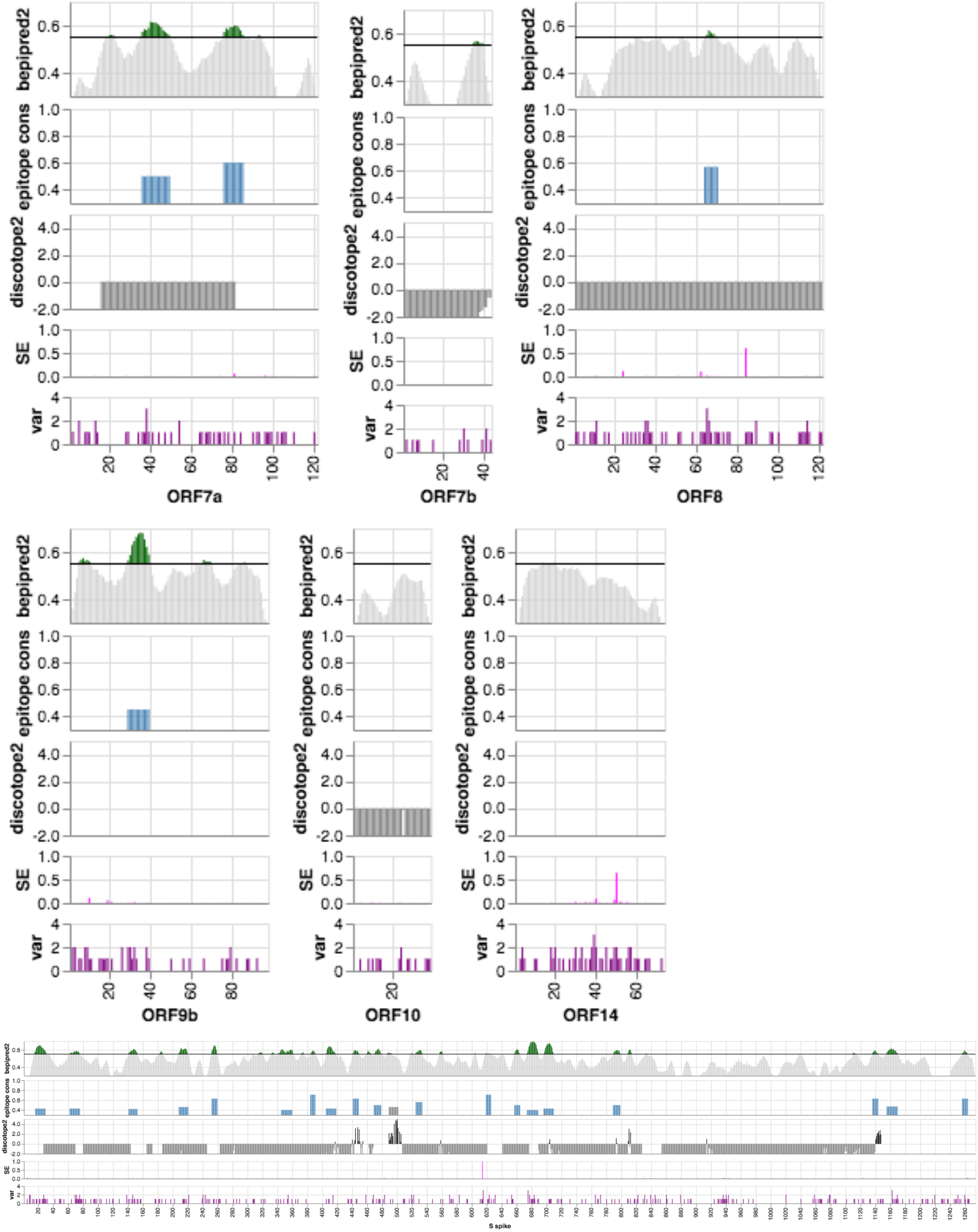
Summary plots for each of the 27 SARS-CoV-2 proteins analyzed in this study. The x-axis represents the amino acid positions in the protein sequence. From top to bottom: **A**) BepiPred2 linear epitope predictions are shown as green bars above the black line representing the 80% specificity threshold (0.55), predictions below the threshold are greyed out; **B**) Epitope conservation vs endemic HCoVs as blue bars spanning the length of the trimmed epitope, with the height representing the degree of conservation (we consider values > 60% as likely non-specific), grey bars correspond to continuous, conformational epitopes with a linear prediction score below the threshold; **C**) DiscoTope2 conformational epitope predictions as black bars above the 80% specificity threshold (−2.5 scaled to zero), predictions below the threshold are greyed out; **D**) Shannon Entropy representing residue variability as pink bars; **E**) Variant counts as purple bars representing the number of residue types found at that position, that differ from the reference sequence.

**Table S2.** Number of protein sequences from HCoV endemic strains found in the public databases. NCBI sequences originate from the Identical Protein Groups (IPG) database and UniProt sequences from the combined UniProtKB databases TrEMBL and Swiss-Prot with VARSPLIC extraction.

| HCoV strain | protein sequence database |  |  |
| --- | --- | --- | --- |
|  | NCBI IPG | UniProtKB | ViPR |
| HKU1 | 259 | 490 | 1,489 |
| 229E | 310 | 457 | 651 |
| OC43 | 1,033 | 1,591 | 5,648 |
| NL63 | 492 | 991 | 1,777 |
| Total | 2,094 | 3,529 | 9,565 |

**Table S3.** Structures and models used for the prediction of conformational epitopes. Min and max positions with structural information are relative to the proteins and may differ from residue numberings in PDB structures and models.

| protein | structure | chain | min | max | source | method | reference |
| --- | --- | --- | --- | --- | --- | --- | --- |
| M membrane protein | C1906TS131_2 | A | 1 | 222 | CASP | de-novo TrRosetta |  |
| N nucleoprotein | 6vyo | A | 50 | 173 | PDB | crystallography |  |
| N nucleoprotein | 6wji | A | 257 | 364 | PDB | crystallography |  |
| ORF3a | C1905TS405_5 | A | 1 | 275 | CASP | de-novo Kiharalab_Z |  |
| ORF6 | C1907TS073_2 | A | 1 | 61 | CASP | de-novo BISCORE |  |
| ORF7a | 6w37 | A | 16 | 81 | PDB | crystallography |  |
| ORF7b | C1910TS131_3 | A | 1 | 43 | CASP | de-novo TrRosetta |  |
| ORF8 | C1908TS131_1 | A | 1 | 121 | CASP | de-novo TrRosetta |  |
| ORF10 | C1909TS210_3 | A | 1 | 38 | CASP | de-novo Kiharalab |  |
| S spike | 6vxx | A | 27 | 1147 | PDB | cryo-EM | Walls 2020 |
| nsp1 | 15661 | A | 12 | 127 | Robetta | homology modelling |  |
| nsp2 | C1901TS156_1 | A | 2 | 639 | CASP | de-novo AlphaFold |  |
| nsp3 PL-PRO | 15659 | A | 1 | 110 | Robetta | homology modelling |  |
| nsp3 PL-PRO | 6vxs | A | 207 | 373 | PDB | crystallography |  |
| nsp3 PL-PRO | 15810 | A | 412 | 540 | Robetta | homology modelling |  |
| nsp3 PL-PRO | 15811 | A | 589 | 676 | Robetta | homology modelling |  |
| nsp3 PL-PRO | 6w9c | A | 748 | 1060 | PDB | crystallography |  |
| nsp3 PL-PRO | 15814 | A | 1091 | 1202 | Robetta | homology modelling |  |
| nsp3 PL-PRO | C1904TS401_1 | A | 1571 | 1927 | CASP | de-novo AlphaFold |  |
| nsp4 | C1902TS210_4 | A | 1 | 500 | CASP | de-novo Kiharalab |  |
| nsp5 3CL-PRO | 6yb7 | A | 1 | 306 | PDB | crystallography |  |
| nsp6 | C1903TS210_4 | A | 1 | 290 | CASP | de-novo Kiharalab |  |
| nsp7 | 7bv2 | C | 2 | 64 | PDB | cryo-EM | Yin 2020 |
| nsp8 | 7bv2 | B | 78 | 191 | PDB | cryo-EM | Yin 2020 |
| nsp9 | 6w4b | A | 5 | 113 | PDB | crystallography |  |
| nsp10 | 6w75 | B | 18 | 139 | PDB | crystallography |  |
| nsp12 RdRp | 7bv2 | A | 32 | 919 | PDB | cryo-EM | Yin 2020 |
| nsp13 helicase | 6jyt | A | 1 | 596 | PDB | crystallography | Jia 2020 |
| nsp14 ExoN-N7-MTase | 15671 | A | 1 | 527 | Robetta | homology modelling |  |
| nsp15 NendoU | 6vww | A | 92 | 346 | PDB | crystallography | Kim 2020 |
| nsp16 MTase | 6w75 | A | 2 | 298 | PDB | crystallography |  |

**Table S4**. Epitope scoring table. <excel spreadsheet>

**Table S5.** ELISA testing results summary (positive defined as values greater than the mean of negative controls added to 3-times the standard deviation of the mean).

| Antigen | # SARS-CoV-2 samples |  | # negative controls |  | Sensitivity | Specificity |
| --- | --- | --- | --- | --- | --- | --- |
|  | tested | positive | tested | positive |  |  |
| S-RBD | 78 | 72 | 22 | 0 | 92.3% | 100% |
| N-Nterm | 78 | 68 | 22 | 0 | 87.2% | 100% |
| Mix, S-RBD + N-Nterm | 80 | 75 | 106 | 3 | 93.8% | 97.2% |

**Table S6.** List of nucleoprotein sequences in the ViPR database from Seattle endemic HCoV strains matching the predicted dominant epitope nprot_10 on the Nucleoprotein dimerization domain. The SARS-CoV-2 epitope sequence and the matching endemic sequence is shown above the table.

| VIPR ID | Isolate |
| --- | --- |
| APU51924 | Human coronavirus OC43 HCoV_OC43/Seattle/USA/SC831/2016 |
| APU51934 | Human coronavirus OC43 HCoV_OC43/Seattle/USA/SC622/2016 |
| APU51935 | Human coronavirus OC43 HCoV_OC43/Seattle/USA/SC9741/2016 |
| ARA15425 | Human coronavirus OC43 HCoV_OC43/Seattle/USA/SC2269/2016 |
| ARK08639 | Human coronavirus OC43 HCoV_OC43/Seattle/USA/SC2924/2015 |
| ARK08655 | Human coronavirus OC43 HCoV_OC43/Seattle/USA/SC2770/2015 |
| ARK08665 | Human coronavirus OC43 HCoV_OC43/Seattle/USA/SC2730/2015 |
| ARK08674 | Human coronavirus OC43 HCoV_OC43/Seattle/USA/SC2476/2015 |
| ARK08683 | Human coronavirus OC43 HCoV_OC43/Seattle/USA/SC2345/2015 |
| ARU07572 | Human coronavirus OC43 HCoV_OC43/Seattle/USA/SC2481/2015 |
| ARU07591 | Human coronavirus OC43 HCoV_OC43/Seattle/USA/SC2854/2015 |
| ARU07598 | Human coronavirus NL63 HCoV_NL63/Seattle/USA/SC2940/2015 |
| ARU07614 | Human coronavirus OC43 HCoV_OC43/Seattle/USA/SC3118/2015 |
| QEG03735 | Human coronavirus NL63 HCoV_NL63/Seattle/USA/SC0179/2018 |
| QEG03744 | Human coronavirus OC43 HCoV_OC43/Seattle/USA/SC0682/2019 |
| QEG03752 | Human coronavirus NL63 HCoV_NL63/Seattle/USA/SC0768/2019 |
| QEG03761 | Human coronavirus OC43 HCoV_OC43/Seattle/USA/SC0810/2019 |
| QEG03771 | Human coronavirus OC43 HCoV_OC43/Seattle/USA/SC0839/2019 |
| QEG03778 | Human coronavirus OC43 HCoV_OC43/Seattle/USA/SC0841/2019 |
| QEG03798 | Human coronavirus OC43 HCoV_OC43/Seattle/USA/SC9430/2018 |
| QEG03805 | Human coronavirus OC43 HCoV_OC43/Seattle/USA/SC9428/2018 |
| QEG03818 | Human coronavirus OC43 HCoV_OC43/Seattle/USA/SC0776/2019 |

**Table S7.** GISAID acknowledgement by authors and centers. <excel spreadsheet>

